# An immature subset of neuroblastoma cells synthesizes retinoic acid and depends on this metabolite

**DOI:** 10.1101/2021.05.18.444639

**Authors:** Tim van Groningen, Camilla U. Niklasson, Alvin Chan, Nurdan Akogul, Ellen M. Westerhout, Kristoffer von Stedingk, Mohamed Hamdi, Linda J. Valentijn, Sofie Mohlin, Peter Stroeken, Nancy E. Hasselt, Franciska Haneveld, Arjan Lakeman, Danny A. Zwijnenburg, Peter van Sluis, Daniel Bexell, Igor Adameyko, Selina Jansky, Frank Westermann, Caroline Wigerup, Sven Påhlman, Jan Koster, Rogier Versteeg, Johan van Nes

## Abstract

Neuroblastoma is a pediatric tumor of the adrenergic sympathetic lineage. Most high risk neuroblastoma go in complete clinical remission by chemotherapy, which is subsequently complemented by retinoic acid (RA) maintenance therapy. However, by unresolved mechanisms most tumors ultimately relapse as therapy-resistant disease. Neuroblastoma cell lines were recently found to include, besides lineage committed adrenergic (ADRN) tumor cells, also immature mesenchymal (MES) tumor cells. Here, we report that MES-type cells synthesize RA and require this metabolite for proliferation and motility. MES cells are even resistant to RA *in vitro*. MES cells appear to resemble Schwann Cell Precursors (SCP), which are motile precursors of the adrenergic lineage. MES and SCP cells express shared RA-synthesis and RA-target genes. Endogenous RA synthesis and RA resistance thus stem from normal programs of lineage precursors that are maintained in an immature tumor cell fraction. These cells are fully malignant in orthotopic patient-derived xenograft models and may mediate development of drug-resistant relapses.

## Introduction

The pediatric tumor neuroblastoma emerges from the peripheral sympathetic nervous lineage. Despite intensive treatment, the outcome for patients with high-stage neuroblastoma remains poor, with an overall survival rate of less than 50%^1,2^. Standard treatment of high-risk neuroblastoma includes several courses of induction chemotherapy, followed by surgical resection of the tumor. Patients are subsequently treated with courses of retinoic acid maintenance therapy and anti-GD2 immunotherapy to eradicate residual tumor cells^3,4^. Most high-stage neuroblastoma respond to these therapeutic treatments by complete clinical remission. However, the majority of them ultimately relapse as drug-resistant and lethal disease.

Retinoic acid (RA) induces differentiation of neuroblastoma cells *in vitro*^5,6^. RA has therefore been included in the treatment of neuroblastoma to promote the differentiation of residual tumor cells and improve outcome. However, the clinical effects of RA on overall survival are modest^4,7^. It is unknown whether neuroblastoma tumors develop resistance to RA *in vivo. In vitro* experiments have identified several genes that can mediate RA resistance in neuroblastoma cell lines^8,9^.

We and others previously showed that neuroblastoma cell lines can be composed of phenotypically divergent cell types^10,11^. ADRN cells are lineage-committed and express transcription factors of the adrenergic lineage, e.g. *PHOX2A, PHOX2B, ASCL1* and *GATA3* as well as enzymes of the catecholamine biosynthesis route, like DBH, DDC and TH. MES-type neuroblastoma cells lack expression of these adrenergic markers but instead express mesenchymal marker genes e.g. *VIM, FN1* and *SNAI2*. MES cells express gene signatures of neural crest cells. MES and ADRN cells each have unique sets of lineage-specific super-enhancers and associated Core Regulatory Circuitries (CRCs), which are assumed to impose lineage identity^12,13^. Indeed, some MES-type CRC transcription factors are able to transdifferentiate ADRN cells into MES cells, including transcriptional and epigenetic reprogramming^10,14^. Phenotypically, MES cells are highly migratory and are *in vitro* resistant to a variety of chemotherapeutic drugs, as compared to ADRN cells. The prevalence of MES-type neuroblastoma cells *in vivo* is less clear. In two reports, no MES-type cells were identified in tumors^15,16^. However, a recent single cell RNAseq analysis identified MES-like primary neuroblastoma that contained tumor cells with features of several developmental cell types from the adrenergic lineage^17,18^. Also several pre-print reports suggest the existence of immature tumor cells in neuroblastoma tumors^19-21^.

Here, we show that undifferentiated MES-type neuroblastoma cells metabolize retinol to produce endogenous RA. These cells use RA to stimulate their proliferation and migration in a gene expression program that mimics developmental stages of the adrenergic lineage.

## Results

### Mesenchymal neuroblastoma cells have an active RA synthesis pathway

As RA is a long standing component of neuroblastoma treatment, we investigated whether it differentially affects ADRN- and MES-type cells. We studied the response to RA in four pairs of isogenic MES and ADRN neuroblastoma cell lines. Each pair has been derived from the tumor of one neuroblastoma patient and consists of >95% homogeneous MES or ADRN cells^10^. In each pair, RA strongly impaired the viability of ADRN-type cells, while MES-type cells were relatively resistant and continued to grow (Fig. 1a and data not shown). Consistently, treatment with RA strongly reduced the S-phase of ADRN-type cells, but MES-type cells largely preserved their S-phase (Supplementary Fig. 1a, b). MES cells are therefore *in vitro* resistant to the differentiating and anti-proliferative effects of exogenous RA. A limited number of cell types can synthesize RA during early embryogenesis^22,23^. We investigated whether MES-type neuroblastoma cells have this property. Endogenous RA synthesis in cells starts with conversion of retinol to retinal by RDH10, followed by the generation of RA from retinal by either ALDH1A3, ALDH1A1 or ALDH3B1^22,24,25^ (Supplementary Fig. 2a). All MES cell lines specifically expressed *ALDH1A3, ALDH1A1* and/or *ALDH3B1* mRNA, while ADRN cell lines hardly expressed these genes (Fig. 1b, c, Supplementary Fig. 2b and Supplementary Table 1). MES-specific expression of ALDH1A3 and ALDH1A1 proteins was confirmed in three isogenic cell line pairs (Fig. 1d and Supplementary Fig. 2c). ChIP-sequencing of H3K27ac showed strong super-enhancers in the vicinity of the *ALDH1A3* and *ALDH3B1* loci, which were strictly associated with expression of these genes, indicating that they belong to the core network of genes with MES-specific super-enhancers^10^ (Fig. 1b and Supplementary Fig. 2b-d).

**Figure 1.**
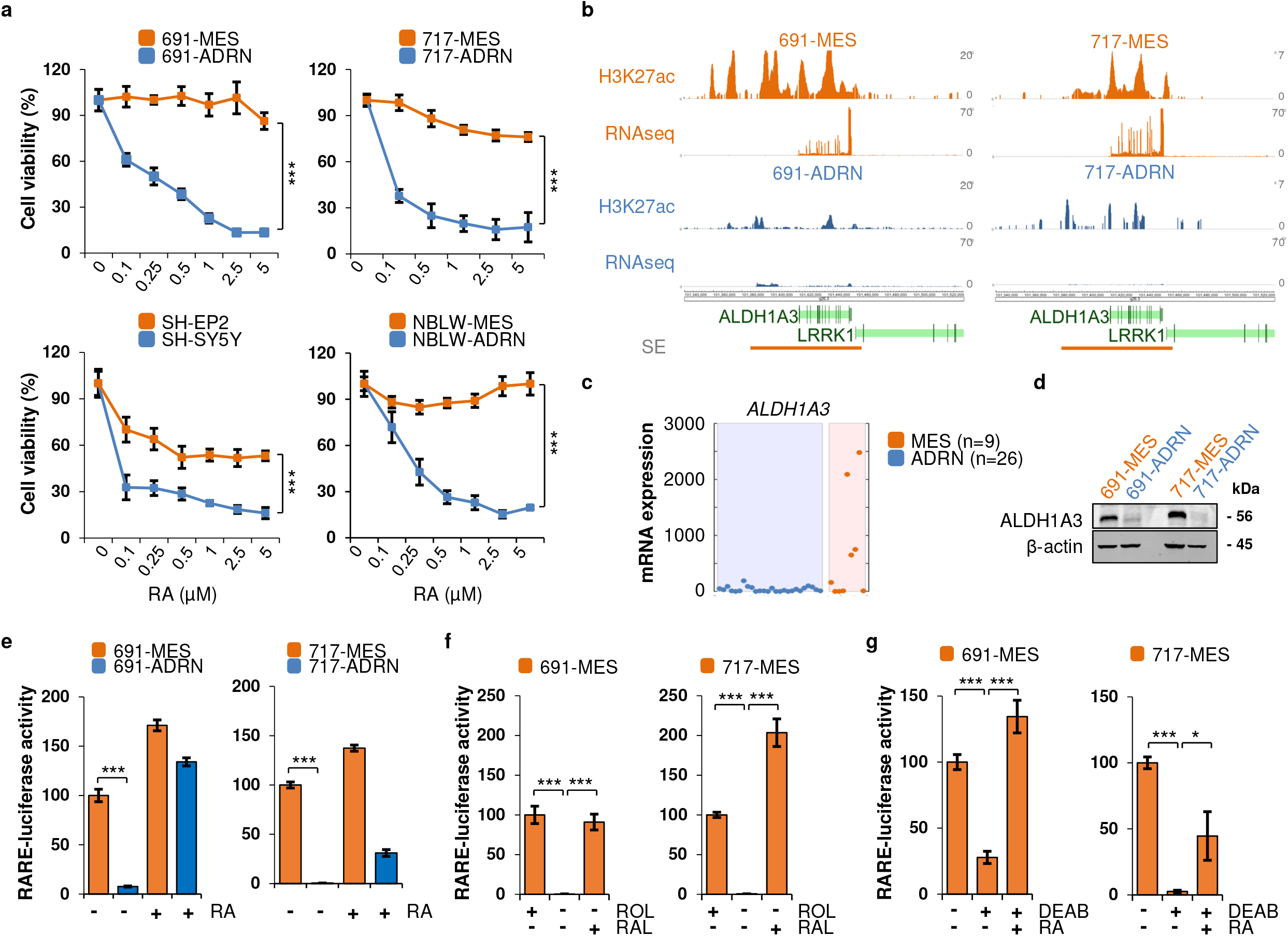
MES-type neuroblastoma cells are resistant to retinoic acid and have an endogenous retinol-to-retinoic acid synthesis pathway. A. CyQuant cell viability assay of four isogenic pairs of MES- and ADRN-type cell lines treated with increasing concentrations of Retinoic Acid (RA). Two-sided Student’s *t*-test assuming equal variance was used to calculate statistical significance, *** *p* < 0.001. MES cells (691-MES, 717-MES, SH-EP2 and NBLW-MES) are depicted in orange, while ADRN cells (691-ADRN, 717-ADRN, SH-SY5Y and NBLW-ADRN) are depicted in blue. B. H3K27ac ChIP-sequencing analysis of the genomic region around the *ALDH1A3* gene spanning positions 101,340,000-101,535,000 on chromosome 15. The y-axis shows reads per 20 million mapped sequences. Lineage-specific super-enhancers of MES cells were identified according to^10^ and are indicated by a horizontal orange bar. RNA sequencing data is shown as reads per 20 million mapped reads and plotted on the y-axis. C. Expression of *ALDH1A3* mRNA measured by Affymetrix gene expression profiling of cell lines of MES (*n* = 9) or ADRN (*n* = 26) phenotype. D. Western blot analysis of ALDH1A3 in 691-MES and -ADRN (left) and 717-MES and - ADRN (right) cells. β-actin is used as loading control. E. RA reporter assay in isogenic MES- and ADRN cell line pairs derived from patients 691 (left) and 717 (right). Endogenous RA reporter activity is measured in the absence (-) of exogenous RA, while an external source of RA (+) transactivates the 3xRARE-luciferase reporter. F. RA reporter assay in 691-MES and 717-MES cells, cultured in the presence (+) or absence (-) of retinol (ROL) or retinal (RAL). G. RA reporter assay in 691-MES and 717-MES cells incubated in the presence (+) or absence (-) of 100 µM of the ALDH inhibitor DEAB or 100 nM RA. The normalized luciferase activities in E-G are ratios between firefly-luciferase values of the 3xRARE reporter and renilla-luciferase values of the transfection control. Error bars denote standard deviation. Two-sided Student’s *t*-test assuming equal variance was used to calculate statistical significance, * *p* < 0.05, *** *p* < 0.001. Source data for A, D and E-G are provided as a Source Data file.

**Figure 2.**
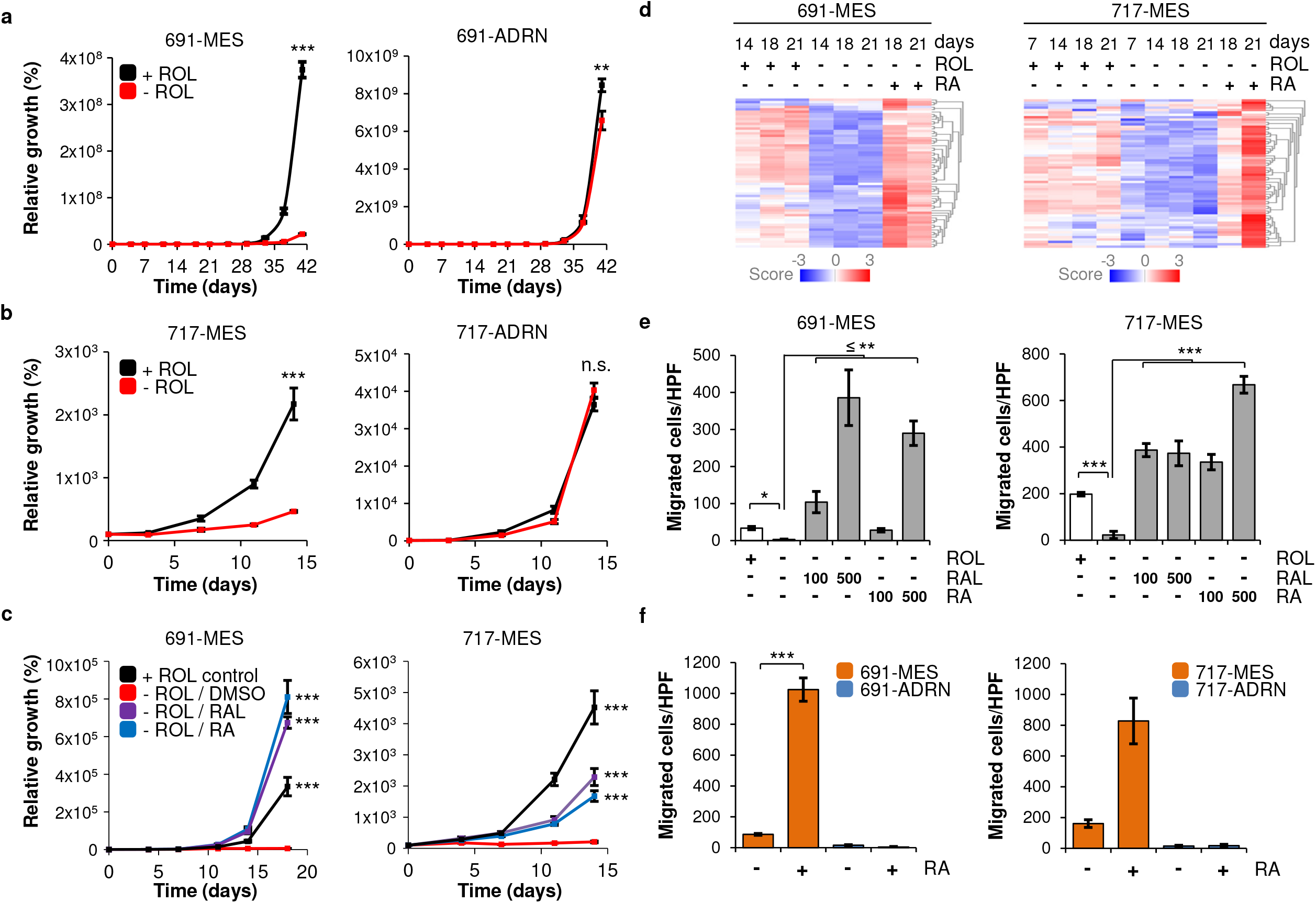
Retinoic acid induces proliferation and migration of MES cells. A, B. Cell count assay (see Methods for description) of (A) 691-MES (left) and 691-ADRN cells (right) and (B) 717-MES (left) and 717-ADRN cells (right) cultured in neural stem cell medium with (+ROL, black) or without retinol (-ROL, red). C. Rescue of 691-MES and 717-MES cells that were cultured in the absence of retinol (-ROL). Cells were pre-cultured without retinol prior to supplementation of the medium with retinal (RAL, 100 nM), retinoic acid (RA, 100 nM) or DMSO at day 0. Proliferation of cells in the presence of retinol (+ ROL) is shown as control. Source data for A-C are provided as a Source Data File. D. Z-score of mRNA expression of RA-induced target genes in MES cell lines. 691-MES and 717-MES were cultured in the presence (+) or absence (-) of retinol (ROL) in a time-course mRNA analysis of three weeks. RA (1 µM) was added from day 18 to day 21 to identify a core set of RA-induced genes in MES cells. The list of RA-target genes in MES cells is available from Supplementary Table 2. E. Transwell migration assay of 691-MES and 717-MES cells in the presence (white bars) or absence (grey bars) of retinol (ROL). Cells were seeded in Boyden chambers in medium supplemented with 100 or 500 nM of retinal (RAL) or RA. Cells were allowed to migrate for 48 hours. Note that 100 nM RA or RAL are sufficient to rescue migration of 691-MES and 717-MES cells to the level of cell migration observed in control cells that are cultured in the presence of ROL. F. Transwell migration assay of MES (orange) or ADRN (blue) cell lines of 691 or 717 in the presence (+) or absence (-) of 1 µM RA. Cells were allowed to migrate for 48 hours. Error bars in all panels depict standard deviation. Two-sided Student’s *t*-test assuming equal variance was used to calculate statistical significance, * *p* < 0.05, ** *p* < 0.01, *** *p* < 0.001. Source data for E, F are provided as a Source Data File.

The Aldefluor assay^26^ showed a high ALDH enzymatic activity in the MES-type cells of the four isogenic cell line pairs, but not in the ADRN-type cells (Supplementary Fig. 3a-e). This activity was blocked by the ALDH-inhibitor DEAB (Supplementary Fig. 3a-d). We therefore studied whether MES cells can synthesize RA. RA binds to multiple RA receptors (RARs) and this complex activates Retinoic Acid Response Elements (RAREs) in gene promoters^22,23^. A RARE-reporter (3xRARE-luciferase) was active in the MES cell lines 691-MES and 717-MES, but not in their isogenic ADRN counterparts (Fig. 1e and Supplementary Dataset 1). To test whether this RARE activity was caused by endogenous RA synthesis, we cultured the MES cells in retinol-deprived medium. This abrogated the RARE-activity, which could be rescued by addition of exogenous retinal (Fig. 1f, Supplementary Fig. 2a and Supplementary Dataset 1). Furthermore, RARE activity of MES cells was blocked by the RARα inhibitor ER50891 and by the pan-RAR inhibitor BMS493 (Supplementary Figs. 2a and 4a-d). Also, inhibition of ALDH by DEAB reduced activity of the RARE-reporter, which was rescued by exogenous RA (Fig. 1g, Supplementary Fig. 2a). Consistently, silencing of ALDH1A3 impaired RA reporter activity, establishing a functional role of ALDH1A3 in the RA pathway (Supplementary Fig. 4e, f). We conclude that MES cells have an endogenous RA synthesis pathway.

**Figure 3.**
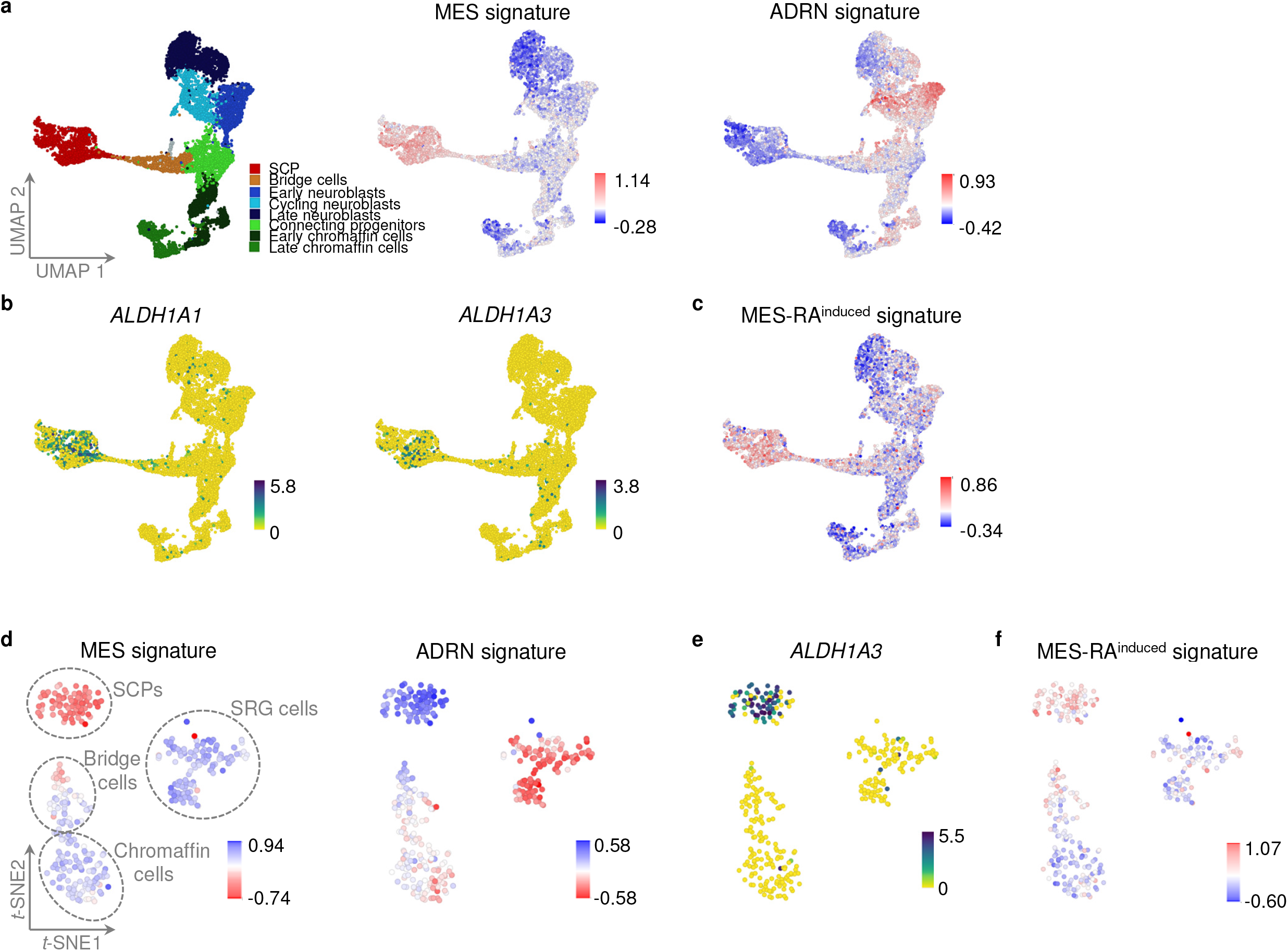
MES-type neuroblastoma cells resemble Schwann Cell Precursors of the developing adrenergic lineage. A. Visualization of gene expression signatures for MES and ADRN neuroblastoma cells^10^ on single-cell analysis of the human adrenal lineage^17^. Cell types of the adrenal lineage are indicated in the left panel. MES and ADRN signatures are indicated in the middle and right panels. The scale indicates the summed z-score of genes from each signature. B. Expression of *ALDH1A1* and *ALDH1A3* mRNA in cell types of the human adrenergic lineage. Color scale shows ^2^log-transformed expression. C. Expression of the MES-specific RA-target gene signature in the human adrenergic lineage. The signature scale shows the summed z-scores of expressed genes of the signature in each cell. D. Gene expression signatures of MES and ADRN neuroblastoma cells^10^ visualized on *t*-SNE analysis of single-cell RNA sequencing analyses of the developing murine adrenergic lineage^29^. The scale indicates the summed z-score of genes from each signature. Schwann Cell Precursors (SCPs), bridge cells, Chromaffin cells and Suprarenal Ganglion (SRG) cells are indicated. Perplexity 12 is chosen for visualization of the *t*-SNE map. E. Expression of *Aldh1a3* mRNA in the murine adrenergic lineage. Color scale shows ^2^log-transformed expression. F. Expression of the MES-specific RA-target gene signature, visualized on single-cells of the developing murine adrenergic lineage. The signature scale shows the summed z-scores of expressed genes of the signature in each cell.

**Figure 4.**
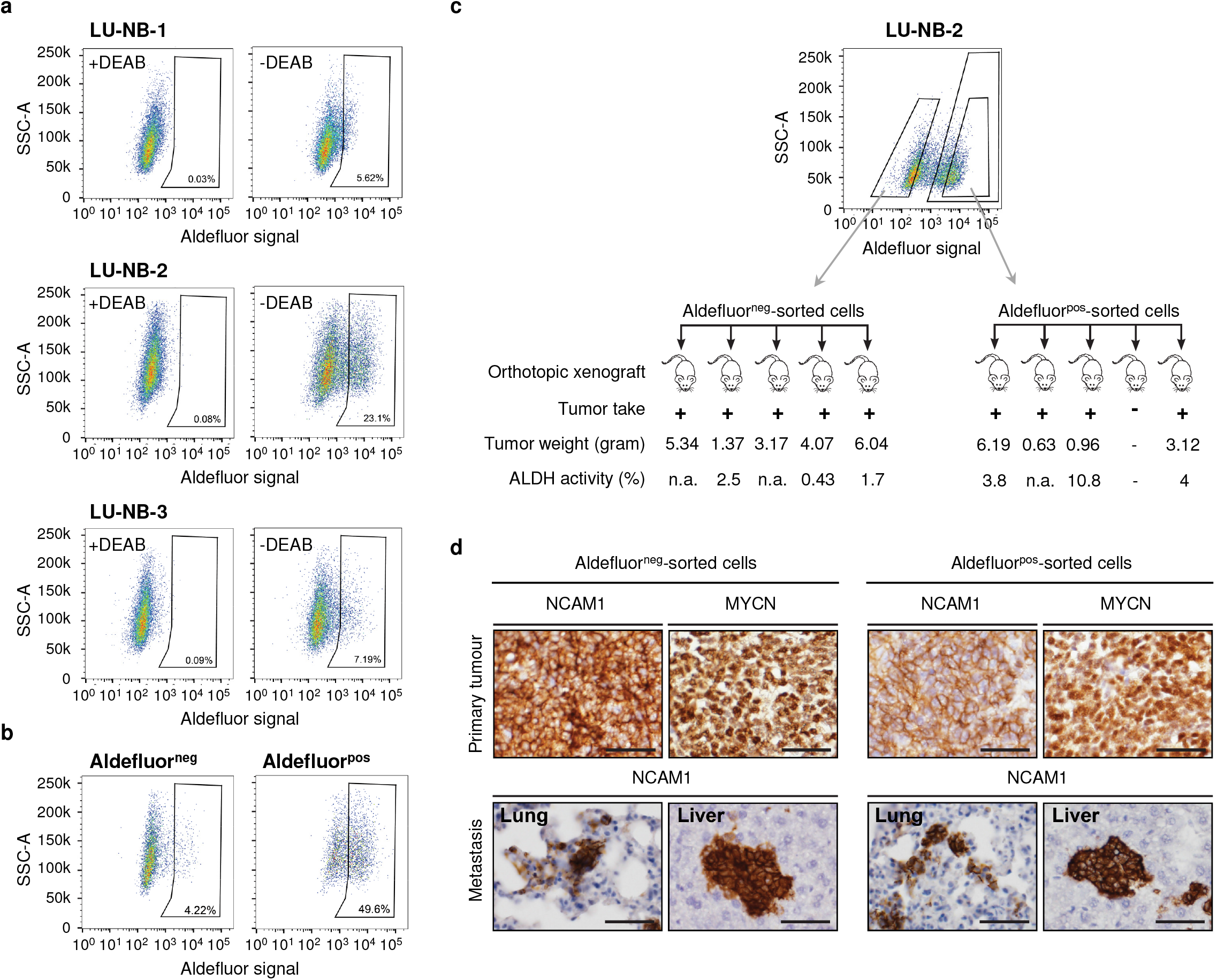
ALDH^positive^ neuroblastoma cells are oncogenic in an orthotopic PDX model and form ALDH^positive^ and ALDH^negative^ heterogeneous tumors. A. Flow cytometry analysis of ALDH activity in cells from the tumors of three orthotopic neuroblastoma PDX tumors (LU-NB-1, LU-NB-2 and LU-NB-3). The corresponding DEAB controls for each sample are shown in the left panels, while the gate marks Aldefluor^positive^ cells in the right panels. Aldefluor-activity and side-scatter (SSC) are shown on the x-axis and y-axis, respectively. B. Interconversion analysis of Aldefluor^positive^ and Aldefluor^negative^ cells. Cells from PDX LU-NB-2 were harvested and sorted in Aldefluor^positive^ and Aldefluor^negative^ cells that were subsequently cultured *in vitro* for T=14 days and re-analysed by FACS for ALDH-activity. The gate indicates ALDH^positive^ cells. Note that Aldefluor^positive^ cells generate Aldefluor^negative^ cells and *vice versa*. Aldefluor-activity is shown on the x-axis, side-scatter (SSC) is shown on the y-axis. C. Aldefluor^positive^ and Aldefluor^negative^ cells were sorted from LU-NB-2 cells and 1×10^4^ cells were orthotopically injected into immunocompromised mice (*n* = 5 per group). At sacrifice, tumor take and tumor weight (in grams) were determined. Cells were isolated from three tumors of each mouse group and analysed for ALDH activity by Aldefluor assay. The percentage of Aldefluor^positive^ cells is indicated. D. Primary tumors as well as the lungs and liver were isolated from all tumor-bearing mice. Stained for NCAM1 and/or MYCN expression revealed metastatic growth in the lungs and livers. Scale bar, 50 µm.

### Mesenchymal neuroblastoma cells require retinoic acid for proliferation and migration

We investigated the functional role of endogenous RA synthesis in MES cells. Blocking of the RA synthesis route by depletion of retinol in the culture medium resulted in decreased proliferation of MES cells, but did not affect ADRN-type cells (Fig. 2a, b). MES cells detached from the culture dish and obtained a sphere-like phenotype (Supplementary Fig. 2e). Addition of retinal or RA rescued all of these effects (Fig. 2c and Supplementary Fig. 2f). Consistently, treatment of two MES and ADRN cell line pairs with DEAB inhibited proliferation of MES cells by ∼50% but did not affect ADRN cells (Supplementary Fig. 2g). RA therefore spurs proliferation of MES cells.

We investigated the target genes of endogenous RA signaling in MES cells by mRNA profiling of retinol-deprived 691-MES and 717-MES cells treated with or without exogenous RA. This identified 62 shared RA-induced genes (logfold 1, *p* < 0.0025), which were enriched for motility and cellular matrix ontologies (Fig. 2d and Supplementary Tables 2 and 3). MES cells are, in contrast to ADRN cells, highly motile^10^ and we therefore tested whether RA supports motility of MES cells. Retinol depletion completely blocked intrinsic motility of MES cells, which was rescued by exogenous retinal or RA (Fig. 2e). The pan-RAR inhibitor BMS493 and the RARα inhibitor ER50891 blocked the intrinsic motility of MES cells in a dose-dependent way (Supplementary Fig. 5a). DEAB treatment of 691-MES and 717-MES inhibited expression of the RA-induced genes and reduced the intrinsic migration of these cells (Supplementary Fig. 5b, c). These data show that endogenous RA controls a gene expression program that confers a migratory phenotype to MES cell lines. Addition of exogenous RA further increased the motility of MES cells, which was blocked by each of the RAR inhibitors (Fig. 2f and Supplementary Fig. 5a). We conclude that endogenous RA signaling is required for the proliferation and migration of MES neuroblastoma cells.

### RA downstream pathways differ in MES and ADRN cells

The downstream pathway of RA signaling identified in MES cells does not explain why MES cells are resistant and ADRN cells are sensitive to RA. As a first analysis of this question, we analyzed the RA response pathway in ADRN-type cells. Exogenous RA did not induce motility of ADRN cells (Fig. 2f). mRNA profiling of 691-ADRN and 717-ADRN identified 98 shared RA-induced genes (≥ logfold 1, *p* < 0.0025). Only 10 of these genes overlapped with the RA-induced genes in MES cells (Supplementary Fig. 6a, b). The RA-target genes in ADRN cells were enriched for neuronal differentiation ontologies (Supplementary Tables 2 and 4). In search for the reason why RA induced different gene sets in MES and ADRN cells, we analyzed whether these genes differ in epigenetic modifications in the different cell types. Analysis of H3K27ac and H3K4me3 ChIP-seq data did not identify clear differences of these marks around the TSSs of the RA-target genes in MES versus ADRN cells (data not shown). The mechanism by which RA activates different gene sets in MES and ADRN cells therefore needs further analyses. The gene sets induced in both cell types are consistent with the finding that RA induces differentiation in ADRN cells, but proliferation and migration in MES cells.

### Heterogeneity of neuroblastoma cells reflect developmental stages of the adrenergic lineage

Neuroblastoma emerges from the peripheral adrenergic lineage^27^. During early embryogenesis, progenitor cells delaminate from the neural crest and migrate ventrally to form the adrenergic lineage. We therefore asked whether the RA-induced motility program of MES cells has an embryonal origin. Single-cell RNA-sequencing of the developing human adrenergic lineage recently characterized three main cell types in this lineage, i.e. Schwann Cell Precursors (SCPs), chromaffin cells and neuroblasts^17,28^. SCPs give rise to chromaffin cells and neuroblasts via populations of bridge cells and progenitor cell states. The clusters of early- and cycling neuroblasts were positive for the previously established ADRN tumor cell signature^10^, but negative for the MES tumor cell signature (Fig. 3a). In contrast, the cluster of SCP cells was strongly positive for the MES signature but negative for the ADRN signature (Fig. 3a). The SCP cells strongly expressed *ALDH1A1 and ALDH1A3*, as well as the signature of the RA-induced genes in MES cells (Fig. 3b,c). This suggests that SCP cells are able to synthesize RA and express RA-induced motility genes, consistent with their migratory phenotype. Similarly, single-cell RNA-sequencing of the mouse adrenergic lineage identified SCPs, chromaffin cells and suprarenal ganglion cells^29^. The mouse suprarenal ganglion cells were positive for the ADRN signature, while SCPs were positive for the neuroblastoma MES signature, expressed *Aldh1a3* and the signature of RA-induced motility genes (Fig. 3d-f). These data suggest a conserved route of RA-synthesis and expression of RA-induced motility genes in SCP cells, consistent with their migratory phenotype. ADRN neuroblastoma cells thus resemble normal neuroblasts, while MES neuroblastoma cells mirror normal SCP cells, providing an explanation for their RA synthesis, motility and RA resistance from a developmental perspective.

### ALDH-positive neuroblastoma cells are malignant

As MES cells are resistant to chemotherapy and RA and may escape standard neuroblastoma therapy, we analyzed the *in vivo* properties of these cells. We investigated a series of 183 stage 4 neuroblastoma^30,31^ for activity of RA pathway genes. The human MES signature showed a gradient of expression in these tumors, in agreement with the described variable proportion of MES-like neuroblastoma cells in tumors^10,11,17^. Human and mouse SCP signatures strongly correlated with the neuroblastoma MES signature and with the signature for RA-induced genes in MES cells (*p* = 1.15×10^−18^ to 6.27×10^−80^, Supplementary Fig. 7a, Supplementary Table 5). Expression of *ALDH1A1, ALDH1A3* and *ALDH3B1* strongly correlated with the neuroblastoma MES signature and the signature for RA-target genes in MES cells (*p* = 1.76×10^−9^ to 9.18×10^−30^, Supplementary Fig. 7b-d and Supplementary Table 5). These analyses are in agreement with RA synthesis capacity of MES cells *in vivo*.

As MES cells might escape current neuroblastoma therapy, we investigated whether this cell population can initiate tumor outgrowth. Three orthotopic patient-derived xenograft (PDX) models of neuroblastoma^32^ were serially passaged. qRT-PCR confirmed *ALDH1A3* expression in cells from all three PDX models (Supplementary Fig. 8a). Dissociation of harvested tumors revealed small subpopulations of cells with ALDH activity (Fig. 4a). Cells from PDX LU-NB-2 were FACS-sorted in ALDH^pos^ and ALDH^neg^ populations. After two weeks of *in vitro* culture, each population had become heterogeneous again, showing spontaneous and bidirectional transdifferentiation of both cell types *in vitro* (Fig. 4b). We transplanted 1×10^4^ sorted ALDH^pos^ or ALDH^neg^ cells in mouse adrenal gland fat pads. After 5 months, ALDH^neg^ cells had formed tumors in 5/5 mice and ALDH^pos^ cells in 4/5 mice (Fig. 4c). Tumors from ALDH^pos^ cells formed metastases in liver and lungs similar to the tumors from ALDH^neg^ cells (Fig. 4d). Analysis of the tumors showed that they had become heterogeneous for ALDH activity (Fig. 4c). We conclude that ALDH^pos^ cells represent a tumorigenic cell population that can recapitulate heterogeneous neuroblastoma.

These results may suggest that ALDH^pos^ and ALDH^neg^ cells in these *in vivo* models would differentially respond to RA. However, RA has been found to be a poor drug in several subcutaneous xenograft models of neuroblastoma^33,34^. We validated these previous observations in xenografts of neuroblastoma cell lines SH-SY5Y and KCNR. These cell lines have an ADRN phenotype *in vitro* and form predominantly ADRN-type tumors (data not shown). Daily treatment with various concentrations of RA as a single drug (2.5, 5, and 10 mg RA per kg body weight per day) did not attenuate tumor growth in both models (Supplementary Fig. 8b-e). Although RA can differentiate ADRN neuroblastoma cells *in vitro*, it is therefore an inefficient drug *in vivo*. It is thus unlikely that RA would promote selective outgrowth of MES-type cells during treatment of primary tumors. Clinically, RA is used in a minimal residual disease setting, following completion of chemotherapy treatment. Further research is needed to answer the question which type of neuroblastoma cells survives chemotherapy and persists during minimal residual disease and how they respond to RA maintenance therapy.

## Discussion

In an increasing number of tumor types, individual tumors appear to include a minor fraction of immature tumor cells that lack lineage differentiation markers^35-44^. The immature cells can be present in treatment-naïve tumors and are often drug-resistant. This has raised the hypothesis that such cells are a source of drug-resistant tumors and relapses. Combination therapy targeting both immature tumor cells and lineage-differentiated tumor cells indeed improved survival in mouse models of several tumor types^35,41,45,46^. Many fundamental questions surround the immature tumor cells, like their origin and the reason of their drug resistance. Here we have addressed the question why immature tumor cells are drug-resistant. RA is used as an anti-cancer drug in several tumor types^47,48^. RA induces differentiation of most neuroblastoma cell lines^5^ and clinical trials showed an improved overall survival in high risk neuroblastoma patients^4^. Nevertheless, the strong *in vitro* effects are thought to translate only in modestly improved clinical outcomes^4,7^.

In isogenic neuroblastoma cell line pairs, we find that RA differentiates ADRN-type cells as expected, but MES-type cells are completely resistant. MES cells even synthesize RA themselves, leading to the paradoxical situation that a subset of tumor cells synthesizes an anti-cancer drug that is intended to kill them. Abrogation of the RA synthesis in MES cells showed that MES cells critically depend on RA for motility and proliferation. RA induced different gene sets in MES cells and ADRN cells. These gene expression programs are associated with neuronal differentiation in ADRN cells, but with motility and migration in MES cells. It is currently not clear why RA induces different gene sets in the two cell types. Thus far, we did not find evidence for a differential epigenetic state of these gene sets in the two cell types, but only a few epigenetic modifications were investigated (H3K27ac and H3K4me3). Apart from differential accessibility of potential RA target genes in MES and ADRN cells, also co-factors might be differentially expressed in both cell types. RA binds to RAR/RXR heterodimers which subsequently activate RARE-elements and transcription of RA-target genes^22,23^. RXR can also form heterodimers with other nuclear receptors^49^. This can lead to cross-regulation between RA, RXR and other nuclear receptor signaling pathways, depending on the dual presence of binding sites for various types of nuclear receptors in regulatory elements^49^. However, none of the various nuclear receptors showed a consistent differential expression in MES and ADRN cells (data not shown). A possible explanation for the observed RA-resistance can be found in various enzymes of the RA-synthesis pathway that control endogenous levels of RA signaling in cells. Co-incidentally, these enzymes of the endogenous RA-dependency pathway may degrade exogenous sources of RA and explain the resistance of MES cells to RA.

It is currently too early to conclude whether, when and where MES-type neuroblastoma cells exist in primary tumors *in vivo*^10,11,15-21^. Analysis of endogenous RA signaling in MES cells may indicate whether these cells are dependent on RA signaling *in vivo*. In addition, this minor population with intrinsic RA dependency and RA resistance can have potential selective advantage during consolidation therapy and may seed relapses.

The RA synthesis pathway also provides insight in the regulatory principles and identity of immature tumor cells. During embryogenesis, immature precursors delaminate from the neural crest and migrate ventrally to target organs where they differentiate. This suggested to us that the motile MES cells relate to migratory precursors of the adrenergic lineage. Development of the human and mouse adrenergic lineage was recently revisited by single cell RNA sequencing^17,28,29^. Three main cell types in the adrenergic lineage were defined at the single cell level: migratory Schwann Cell Precursors (SCP), Chromaffin cells and neuroblasts (referred to as Suprarenal Ganglion cells in mice), which are connected via several intermediate cell states. Here, we found that MES cells resemble SCP cells and reiterate developmental programs from these precursor cells. SCP cells are strongly positive for the previously established signature of MES-specific genes^10^. In contrast, the human ADRN signature was strongly expressed by neuroblasts, which is in line with the presumed origin of neuroblastoma from immature neuroblasts^17,27,28^. SCP cells express the key RA synthesis genes *ALDH1A1* and *ALDH1A3* as well as the signature of RA-induced genes in MES cells. Interestingly, we note that *ALDH1A3* is specifically expressed in early SCPs, while *ALDH1A1* is expressed by early and late SCPs. These developmental differences in timing and expression may explain the patterns of *ALDH1* isoforms in various MES neuroblastoma cell lines. Together, this suggests that MES cells are not simply tumor cells that are de-differentiated and have lost lineage markers, but in fact resemble a specific precursor cell type of the adrenergic lineage. The properties and metabolism of these precursors are faithfully conserved in the MES-type tumor cells. MES and ADRN tumor cells can bi-directionally transdifferentiate into one another^10^. Their resemblance to SCPs and neuroblasts respectively, may represent a differentiation trajectory of the adrenergic lineage.

We previously established that MES cells are chemo-resistant relative to ADRN cells^10^. Here we show that MES cells are also resistant to RA and moreover, that this cell type can grow out to heterogeneous neuroblastoma *in vivo*. The characteristics of MES and SCP cells may lead to the identification of drugs that specifically kill MES-type neuroblastoma cells and can be used to abate the emergence of resistant tumors and relapses.

## Methods

### Cell culture, metabolites and inhibitors

Cell lines SH-SY5Y and SH-EP2 were cultured as described previously^50^. The isogenic cell line pairs from the tumor of patient 691 (691-MES and 691-ADRN), or from the tumor of patient 717 (717-MES and 717-ADRN) were derived and cultured in neural stem cell (NSC) medium as described^10,51^. Retinol-free and retinol-containing medium were prepared by addition of respectively retinol-free B27 (17504-044, Life Technologies) or retinol-containing B27 (12587-010, Life Technologies) to NSC-medium. The NBLW-MES and NBLW-ADRN cell lines were derived from the parental NBLW cell line^52^ and will be described elsewhere (Westerhout, Hamdi et al., ms. submitted for publication.). NBLW, NBLW-MES and NBLW-ADRN cells were cultured in RPMI-1640 medium supplemented with 10% Foetal Calf Serum, 1x Non-essential amino acids, 20 mM L-Glutamine, 10 units/mL penicillin and 10 μg/mL streptomycin (Life Technologies). Cell line identities were verified by short tandem repeat (STR) analysis. Cell lines were routinely checked for the presence of mycoplasma using the MycoAlert detection kit (Lonza). Neuroblastoma patient-derived xenografts (PDXs) were established and maintained as previously described^32^. PDX cell lines were cultured according to^53^ and authenticated by SNP profiling (Multiplexion, Germany). All-trans retinoic acid (RA, R2625), all-trans retinal (RAL, R2500) and diethylaminobenzaldehyde (DEAB, D86256) were from Sigma. The pan-RAR inhibitor BMS493 (3509) and the RARα inhibitor ER50891 (3823) were from Tocris.

### Retinoic acid reporter assay

An RA reporter gene containing a multimerized Retinoic Acid Response Element (3xRARE) upstream of a firefly-luciferase gene^8,9^ was co-transfected with a renilla-luciferase gene in MES (691-MES, 717-MES) and ADRN (691-ADRN, 717-ADRN) cells. DNA (500 ng of 3xRARE-luciferase and 500 ng of renilla luciferase) was co-transfected using FuGENE HD reagent (E2312, Promega). At 24 hours post-transfection, the culture medium was replaced. Lysates for luciferase analysis were harvested at 48 hours after transfection. 100 nM RA and/or indicated concentrations of ER50891 and BMS493 were added to the culture medium 24 hours prior to the harvest of lysates for luciferase analysis. To test the effect of retinol on RA-reporter activity, 691-MES or 717-MES cells were pre-cultured for three days in the presence (+) or absence (-) of retinol prior to transfection of the 3xRARE-luciferease reporter gene. The next day, culture medium (+/-ROL) was replaced and supplemented with 100 nM RAL as indicated, followed by 24 hours culture prior to luciferase analysis. Firefly- and renilla-luciferase activity was determined using the Dual-Luciferase reporter assay system (E1910, Promega) and measured on a Synergy HT microplate reader (BIOTEK). For each sample, the luciferase value was divided by the renilla value to obtain a RARE-activity measurement that is corrected for transfection efficiency. Source data of 3xRARE-luciferase experiments are provided as a Source Data file

### Gene expression profiling and analysis of micro-array data

Total RNA was isolated using Trizol reagent (Invitrogen) and extracted using the RNeasy Mini Kit (Qiagen) according to the manufacturer’s instructions. RNA quality was verified on a Bioanalyzer (Agilent). RNA was hybridized on Affymetrix HG U133A plus2.0 gene chips and normalized using the MASS5.0 algorithm. For RNA-sequencing, libraries were generated using the Kapa RNA HyperPrep kit with RiboErase (HMR, Kapa Biosystems), according to the manufacturer’s instructions. 250 ng of RNA isolated from cell lines 691-MES, 691-ADRN, 717-MES, or 717-ADRN was used as an input for library preparation with 10 cycles of amplification. Libraries were sequenced on a HiSeq4000 (Illumina) with 50 base-pairs single-end reads.

The ADRN-RA gene expression signature was generated from the overlap of regulated genes (≥ logfold 1) in 691-ADRN and 717-ADRN cells treated with 1 µM RA or DMSO as a control and analyzed at 0, 24, 48 or 72 hours of treatment. The MES-RA signature was generated from the overlap of RA regulated genes in 691-MES and 717-MES cells that were cultured in retinol-free medium. First, cell lines were switched from retinol-containing medium to retinol-free culture medium and total RNA was harvested after culturing cells for 0, 14, 18 and 21 days in or retinol-free medium. From day 14 onwards, 1 µM RA was added to rescue gene expression in retinol-deprived cell cultures. RA target genes met the following requirements: ≥ 1 logfold regulated, a minimum of 1 present call, minimum expression of 50 units and P ≤ 0.0025 after RA treatment. Genes regulated in a DMSO control experiment were excluded. Expression data is available from GEO (GSE124960). A MES-RA gene signature score was calculated using a previously described methodology^10^. MES signature genes^10^ were removed from the MES-RA signature score to exclude bias in correlation analyses of these signatures in neuroblastoma tumors.

Expression profiles of MES- and ADRN neuroblastoma cell line pairs are available from GEO (GSE28019 and GSE125059). The cohort of primary human neuroblastoma was described previously and is available from GEO (GSE62564^31^). Bioinformatic analyses were performed using the R2 platform (<u>r2.amc.nl</u>). Gene ontology analysis was performed in R2 using significant (*p* ≤ 0.05) categories in biological processes called between level 3 and 9.

### Single-cell analysis of adrenergic lineage development

Single-cell RNA sequencing of the human adrenergic lineage was recently published^17^. Processed data, as well as their UMAP embedding as used in the publication^17^ was downloaded from (https://adrenal.kitz-heidelberg.de/developmental_programs_NB_viz/). Data was converted into a suitable format for R2 and provided as an interactive UMAP (r2.amc.nl). Single-cell RNA-sequencing of the mouse adrenergic lineage was published^29^ and is available from GEO (GSE99933). For extensive descriptions of single-cell RNA sequencing methodology and filtering steps, we refer to the detailed methods sections of these studies^17,29^. The *t*-distributed stochastic neighbor embedding (*t*-SNE) analysis was performed in the R2 platform (<u>r2.amc.nl</u>). As data, the count information from GSE99933 at NCBI GEO^29^ was used and pre-processed such that every cell was normalized to a count, where the total sum of signal equals to 100000. Every signal was elevated by 1 unit to enable log transformation. This dataset is accessible in R2 as (‘Normal Peripheral Glial Cells E13.5 - Furlan - 376 - custom - gse99933’). The *t*-SNE analysis was performed in R2 on ^2^log transformed zscore values for those genes that had a readcount signal in at least 1 cell using the Rtsne package, with perplexities ranging from 5-50. The resulting 2-dimensional coordinates from perplexity 12 were chosen for visualization and can also be found in R2 (*t*-SNE maps).

The mouse SCP signature was derived from a comparison of differential gene expression (ANOVA on ^2^log transformed values, with FDR-correction), between groups of SCP cells and SRG cells. Only genes with a SCP-specific expression (*r* ≥ 0.7, *n* = 238 genes) were included. For analysis of this SCP signature in human neuroblastoma, human genes were translated to mouse orthologues. For correlation analysis of the SCP signature with the MES gene signature or with the RA^induced^ gene signature, overlapping genes were removed to avoid bias in correlation.

### ChIP-sequencing

ChIP-sequencing was essentially performed as described^10^. Histone-bound DNA from isogenic cell line pairs (691-MES, 691-ADRN, 717-MES, 717-ADRN, NBLW-MES and NBLW-ADRN cells) was precipitated using antibodies against H3K27ac (ab4729, Abcam). ChIP-sequencing of H3K27ac from NBLW-MES and NBLW-ADRN cells and is available from GEO (GSE125059). ChIP-sequencing profiles of H3K27ac in 691-MES, 691-ADRN, 717-MES, 717-ADRN, SH-EP2 and SH-SY5Y cell lines were generated previously^10^ and are available from GEO (GSE90805).

### Transwell migration

Transwell migration assays were performed in ThinCert 24-well transwell inserts (8 μm pore size, 662638, Greiner). 2.5×10^5^ 691-MES or 691-ADRN or 1×10^5^ 717-MES or 717-ADRN cells were allowed to migrate for 48 hours through a transwell to a gradient of B27 (20% B27 in the upper chamber of transwell to 100% B27 as chemoattractant in the lower chamber of 24-well plate) in the presence or absence of retinol. Migration assays were performed in the presence or absence of RA, RAL, DEAB, pan-RAR inhibitor BMS493, or RARα inhibitor ER50891. For retinol deprivation experiments, cells were cultured in retinol-free NSC medium for two weeks prior to seeding in a transwell. To rescue migration of retinol-deprived cell cultures, cells were pre-incubated with RAL or RA for four days prior to seeding in a transwell. Aldehyde function in cell migration was studied by pre-treatment of cells with indicated concentrations of DEAB for four days, prior to cell seeding in transwells in the presence or absence of DEAB. Non-migrated cells were removed using cotton swabs and transwells were washed in PBS. Migrated cells were fixed with 4% PFA (4078-9001, Klinipath) for 10 min., followed by fixation with 50% methanol in PBS for 5 min. and 100% methanol for 20 min. Fixed cells were stained in 0.1% crystal violet.

### CyQuant proliferation assay

Cells were seeded in 96-well plates and treated with indicated concentrations of RA or DEAB. After 5 or 6 days of treatment, DNA content was measured using the CyQuant assay (C35012, Life Technologies) according to the manufacturer’s instructions with the exception that CyQuant reagents were added at half of the indicated volumes. DNA content was measured on a Synergy HT microplate reader (BIOTEK).

### EdU incorporation assay

Cells were seeded in 6 cm dishes and treated with 1, 5 and 10 μM RA for 5 days. 691-MES and 691-ADRN cells received a 2 hour pulse with a final concentration of 10 μM EdU, while 717-MES and 717-ADRN cells were treated for 3 hours with a final concentration of 10 μM EdU. EdU and PI staining were performed using Click-iT Plus EdU Alexa Fluor 488 Flow Cytometry Assay (C10632, Thermofischer Scientifc) according to the manufacturer’s instructions.

### Cell count assays

To determine the effect of retinol deprivation on proliferation of isogenic cell-line pairs, 5×10^5^ 691-MES cells, 5×10^5^ 691-ADRN cells or 4×10^5^ 717-MES cells were seeded in 6 cm^2^ dishes and 4×10^5^ 717-ADRN cells are seeded in 25 cm^2^ flasks in NSC medium supplemented with retinol-containing B27 or retinol-free B27. In this cell count assay, cells were counted and reseeded every week at day 3 and at day 7 for the duration of the experiment, at similar numbers as at the start of the experiment. Cell number was determined using a Coulter Counter (Beckman). To rescue the growth arrest induced by retinol deprivation, 717-MES and 691-MES were pre-cultured in retinol-free medium for 2 or 3 weeks respectively to induce a proliferation phenotype before start of experiment. Subsequently, 4×10^5^ 717-MES cells or 5×10^5^ 691-MES cells cultured retinol-free medium were seeded in 6 cm^2^ dishes in the absence or presence of 100 nM RA or 100 nM RAL. Total cell number was quantified using a coulter counter and cells were reseeded at similar cell densities as at start of experiment every 3^rd^ or 4^th^ day until the end of experiment.

### Quantitative real-time PCR

Total RNA was extracted using the RNeasy Mini Kit (Invitrogen). qRT-PCR for *ALDH1A1* and *ALDH1A3* was performed as described previously^54^. The relative gene expression was normalized to the expression of three reference genes (*SDHA, UBC, YWHAZ*) using the comparative Ct method^31^. Forward (F) and reverse (R) oligo sequences were ALDH1A1-F (5’-TGTTAGCTGATGCCGACTTG-3’), ALDH1A1-R 5’-TTCTTAGCCCGCTCAACACT-3’) and ALDH1A3-F (5’-TCTCGACAAAGCCCTGAAGT-3’, ALDH1A3-R 5’-TATTCGGCCAAAGCGTATTC-3’), SDHA-F (5’-TGGGAACAAGAGGGCATCTG-3’), SDHA-R (5’-CCACCACTGCATCAAATTCATG-3’), UBC-F (5’-ATTTGGGTCGCGGTTCTTG-3’), UBC-R (5’-TGCCTTGACATTCTCGATGGT-3’), YWHAZ-F (5’-ACTTTTGGTACATTGTGGCTTCAA-3’), YWHAZ-R 5’-CCGCCAGGACAAACCAGTAT-3’). Each experimental condition was performed in triplicate.

### Western blot analysis

Total cell lysates were made in RIPA-buffer supplemented with Protease inhibitor cocktail (11836170001, Roche), 1mM NaF and 1mM NaVO3. Western blotting was performed according to standard protocols. In short, protein was transferred to nitrocellulose membrane (GE healthcare, RPN203D). Membranes were blocked for 1 hour at RT, incubated at 4°C overnight with primary antibody (1:1000) and incubated for 1 hour at RT with secondary antibodies in either 2% PBA (GE healthcare, RPN418), 5% ELK or OBB (LI-COR, 829-31080) in PBS with 0.1% TWEEN (Sigma, P1379). Primary antibodies for western blotting were YAP/TAZ (8418, Cell Signaling), ALDH1A1 (54135, Cell Signaling), GATA3 (5852, Cell Signaling), total AKT (4691, Cell Signaling) and ALDH1A3 (ab129815, Abcam). Secondary antibodies for chemiluminescence detection were donkey anti-rabbit-HRP (GE healthcare, NA 9340V, 1:5000) or sheep anti-mouse-HRP (GE healthcare, NXA931, 1:5000). Chemiluminescence detection was performed using the ECL Prime Western Blotting kit (GE-healthcare, RPN2232) and developed on a ImageQuant LAS 4000 (GE healthcare, 28-9558-10). Secondary antibodies for infrared fluorescence detection were donkey anti-rabbit-IRDye® 800CW (Rockland, 611-731-127, 1:5000). For infrared fluorescent detection, membranes were scanned on an Odyssey Infrared imaging System (LI-COR, LIC-9201-00).

### Aldefluor assay

ALDH enzymatic activity was determined using the ALDEFLUOR^™^ kit (#01700, Stem Cell Technologies). For PDX derived tumors and cell-lines, 1×10^6^ cells and for isogenic neuroblastoma cell-line pairs, 5×10^5^ cells were used. Cells were re-suspended in ALDEFLUOR assay buffer and ALDH substrate was added according to manufacturer’s protocol. Half of the cell suspension mixture was immediately mixed with DEAB. Following incubation at 37°C, cells were suspended in ALDEFLUOR assay buffer. PDX derived cells were centrifuged and suspended in ALDEFLUOR assay buffer containing Fixable Viability Stain 660 (FVS660, 1:1000, BD Horizon) after incubation at 37°C. FACS data was acquired on either a FACSVerse instrument (BD bioscience) or Accuri C6 (BD bioscience) and subsequent data analysis was performed using FlowJo software (FlowJo, LLC) or Accuri C6 (BD bioscience) software.

### Animal procedures and immunohistochemistry

Orthotopic injections of PDX cells were performed as previously described^55^. Four-to six-week-old female or male NSG mice were purchased from Charles River (Charles River Laboratories). Mice were housed under pathogen-free conditions. For orthotopic injections of ALDH-positive and ALDH-negative populations, cells were prepared by the ALDEFLUOR^™^ assay (Stem Cell Technologies) and sorted on a FACSAria IIu or FACSAria III instrument using the DIVA software. Live gating was performed using DAPI as a live/dead stain. A small aliquot of sorted populations was re-analyzed directly after sorting to verify the sorting procedure. All animal procedures followed the guidelines set by the Malmö-Lund Ethical Committee for the use of laboratory animals and were conducted in accordance with European Union directive on the subject of animal rights. Experimental protocols were approved by the Malmö-Lund Ethical Committee (ethical permits M146-13 and M11-15).

Xenograft tumors and mice organs were fixed in formalin and embedded in paraffin. After antigen retrieval using PT Link (Dako), 4 µm tissue sections were stained and developed using AutostainerPlus (Dako). Antibodies were diluted in block solution and sections were incubated for 30 minutes with primary antibody and 20 minutes with secondary antibodies. The following antibodies were used: NCAM (Leica Biosystems, NCL-L-CD56-504, 1:50) and MYCN (Novus Biologicals, 23960002, 1:300). Images were acquired using an Olympus BX63 microscope and DP80 camera along with the CellSense Dimension imaging software.

For RA-treatment in neuroblastoma xenografts, SH-SY5Y or KCNR cells were implanted subcutaneously in NMRI-Foxn1^nu/nu^ mice (Charles Rivers; females, 6-8 weeks). RA-treatment (2.5, 5, 10 mg/kg/day or vehicle control) was started at a tumor volume of 125-200 mm^3^. RA was administered intra-peritoneally for five consecutive days, followed by two days off treatment, for a maximum duration of three weeks. The tumor volume was measured twice weekly by a caliper. Mice were sacrificed at the humane endpoint (tumor size > 1200 mm^3^) and the tumors were isolated, fixed in 4% (w/v) buffered formaldehyde (Klinipath) and embedded in paraffin for histological analyses. All animal experiments were conducted under institutional guidelines and according to the law, approved in DAG203AC by the AMC animal ethics committee.

### Statistical analysis

All experimental values are reported as mean ± standard deviation from at least a group size of n=3, unless otherwise stated. A two-sided unpaired Student’s *t* test was used for statistical analyses. The significance of correlations of signatures in tumor series is determined by t=R/sqrt((1-r∼2)/(n-2)), where R is the correlation value and n is the number of samples. Distribution measure is approximately as t with n-2 degrees of freedom.

### Data availability

ChIP-sequencing data, mRNA expression profiles and single-cell expression data used in this study are available from the Gene Expression Omnibus (GEO) with accession numbers GSE124960, GSE90805^10^, GSE28019^10^, GSE125059, GSE62564^31^, GSE99933^29^. Raw data of RARE-luciferase experiments is available for Figs. 1 and Supplementary Fig. 3 and is provided as a source data file. Reagents used in this manuscript are available from the corresponding authors upon reasonable request.

## Supporting information

BioRxiv - van Groningen, Niklasson et al - Supplementary Figures

## Acknowledgments

We thank Sue Cohn for providing the NBLW cell line, Javanshir Esfandyari for help with mouse experiments and Richard Volckmann for processing of bio-informatics data. The RARE-reporter construct was a kind gift of René Bernards. This research was supported by grants from Villa Joep, the European Research Council (ERC-Advanced Grant no. 340735 to R.V.), KiKa (projects 11, 33, 66), Dutch Cancer Society KWF (UVA 2010-4878) and NWO (EraCoSysMed 9003035006), the Swedish Cancer Society (170276, CAN 2017/376, CAN 2017/996), the Swedish Research Council (2017-01304), the Swedish Childhood Cancer Fund (PR2015-0081, PR2017/0003) and Fru Berta Kamprad’s Foundation (BKS 47/2014).

## Author contributions

T.v.G., R. V. and J.v.N. conceived the study, analyzed the data and wrote the manuscript. T.v.G., C.U.N., A.C., N.A., E.M.W., K.v.S., S.M., D.B., C.W. and J.v.N. performed the experiments and analyses. M.H., L.J.V., P.S., N.E.H., F.H., A.L. and P.v.S contributed to experiments. J.K. and D.A.Z. performed bio-informatics analyses. C.U.N., D.B., C.W. and S.P. designed and analyzed experiments, under the supervision of S.P. I.A., S.J and F.W. contributed single-cell expression data. R.V. and J.v.N. supervised the study.

## Additional information

**Supplementary information** is available for this paper.

**Source Data** is available for this paper.

