## Supplementary material for "An immature subset of neuroblastoma cells synthesizes retinoic acid and depends on this metabolite": BioRxiv - van Groningen, Niklasson et al - Supplementary Figures

### Supplementary Figure 1

**a**

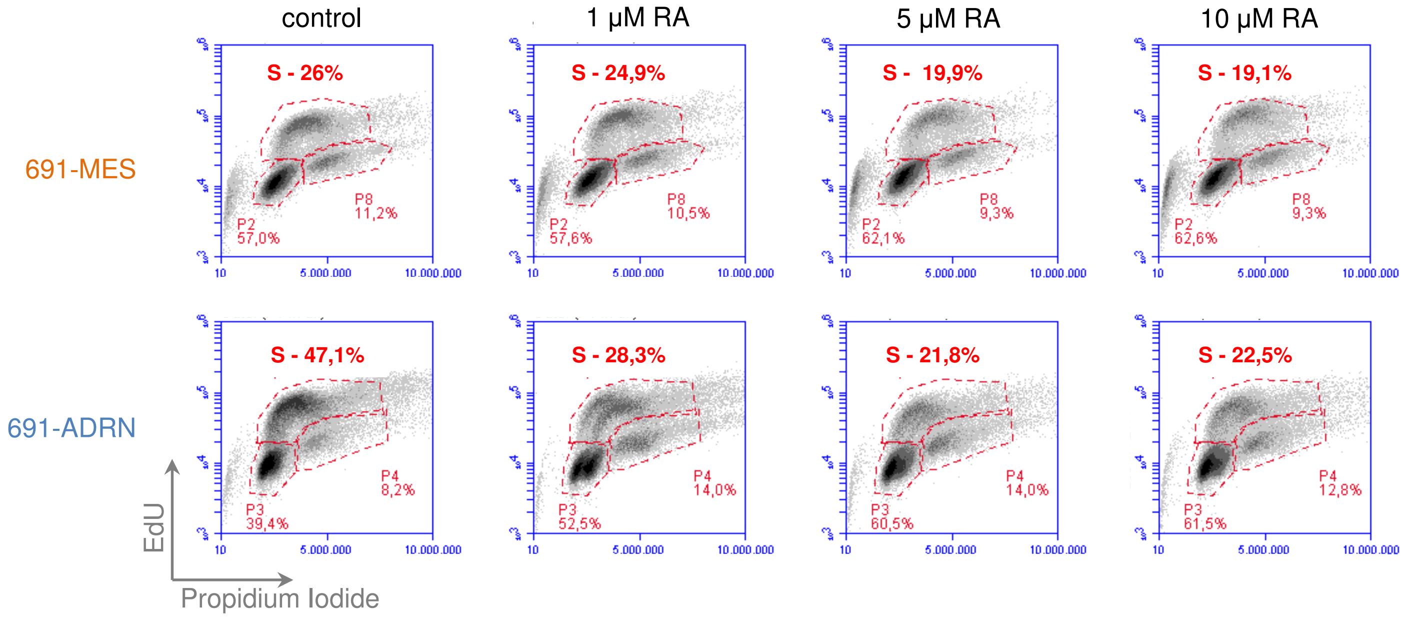


**b**

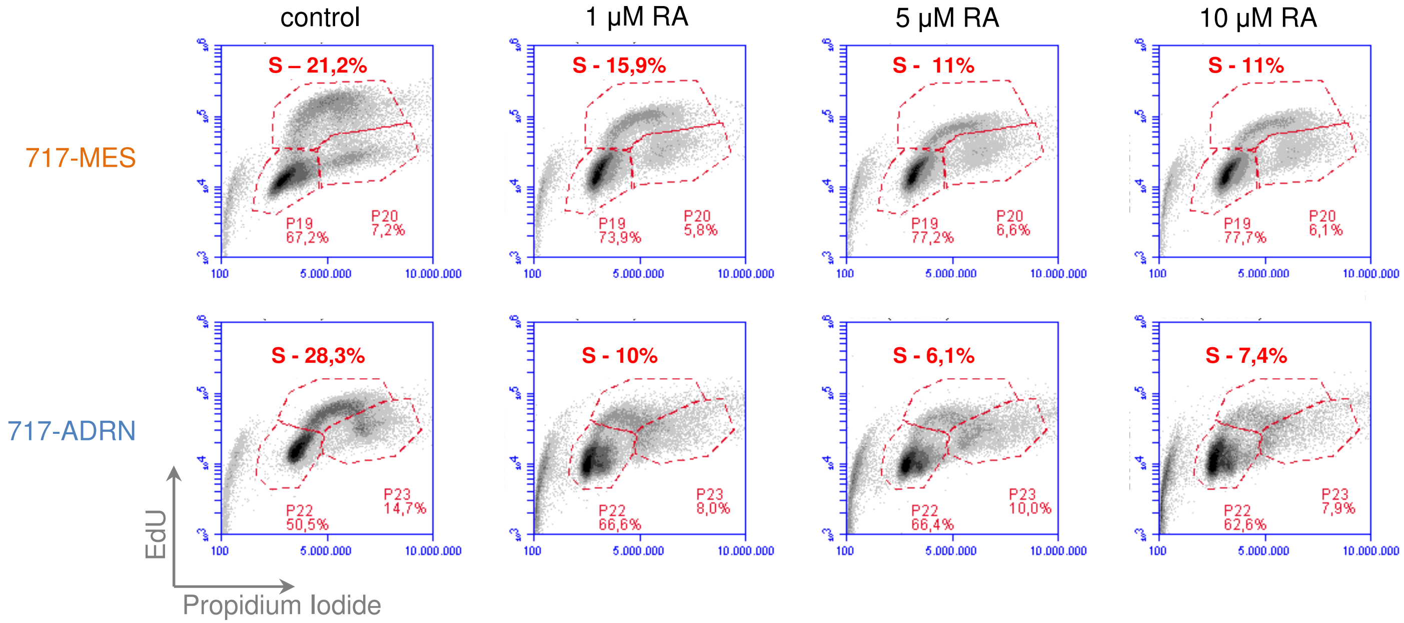

**Supplementary Figure 1 – related to Figure 1.**

**Proliferation of MES and ADRN cells in response to RA.**

A, B. EdU incorporation assay and Propidium-Iodide (PI)-staining for MES and ADRN cells from patient 691 (in A) and patient 717 (in B). 691-MES, 691-ADRN, 717-MES and 717-ADRN cells were treated with 1, 5 and 10 μM RA for 5 days and analyzed by FACS for PI (x-axis) and EdU (y-axis).

**Supplementary Figure 2**

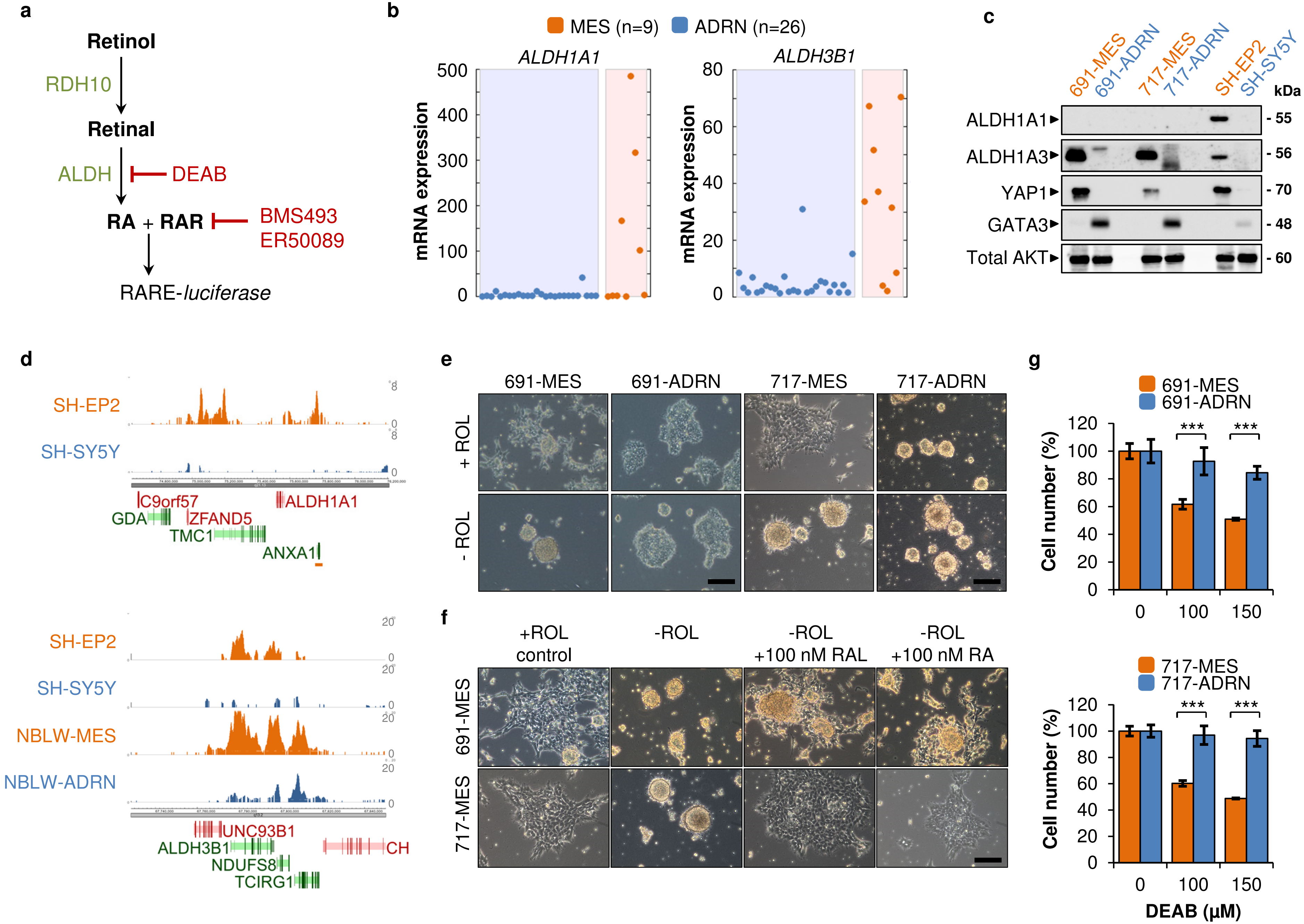

**Supplementary Figure 2 – related to Figure 1.**

**MES neuroblastoma cells have an active RA synthesis pathway.**

A. Schematic overview of the core RA synthesis pathway, according to[15,16]. Retinol is converted to retinal by RDH10. Retinal is converted to RA by ALDH enzymatic activity, the latter can be inhibited by DEAB. RA associates with Retinoic Acid Receptors (RARα, RARβ, RARγ). The RARα inhibitor ER50891 and the pan-RAR inhibitor BMS493 can block the RA/RAR transcriptional activity. A RARE-luciferase reporter gene responds to an endogenous source of RA.

B. mRNA expression of *ALDH1A1* (left) and *ALDH3B1* (right) genes, measured by Affymetrix profiling in MES ($n$ = 9, in orange) and ADRN ($n$ = 26, in blue) cell lines.

C. Western blot analysis of ALDH1A1, ALDH1A3, the MES-marker YAP1 and the ADRN-marker GATA3 in isogenic cell line pairs 691-MES and 691-ADRN, 717-MES and 717-ADRN and SH-EP2 and SH-SY5Y. Cell lines of MES or ADRN phenotype are shown in orange and blue, respectively. Total AKT is shown as loading control.

D. ChIP-sequencing analyses of H3K27ac of the genomic regions around the *ALDH1A1* or *ALDH3B1* genes. The y-axis represents reads per 20 million mapped sequences. The genomic position of the genes on sense (green) or antisense (red) DNA strands is shown on the x-axis.

E. Bright field images of 691-MES, 691-ADRN, 717-MES and 717-ADRN cultured in neural stem cell medium with retinol (+ROL, upper panels) or without retinol (-ROL, lower panels) for 14 days. Scale bars represent 50 µm.

F. Bright field images of 691-MES (upper panels) and 717-MES (lower panels) cells cultured in the presence (+) or absence (-) of retinol (ROL) for 18 days. At day 14, 100 nM Retinal (RAL) or 100 nM RA was added as indicated. DMSO was added as control. Scale bar represents 50 µm.

G. CyQuant cell viability assay of isogenic cell-line pairs 691-MES/-ADRN (left) and 717-MES/-ADRN (right) with increasing concentrations of the ALDH-inhibitor DEAB. MES cell-lines are shown in orange and ADRN cell-lines in blue. Two-sided Student's $t$-test assuming equal variance was used to calculate statistical significance, *** $p < 0.001$. Source data are provided as a Source Data File.

#### Supplementary Figure 3

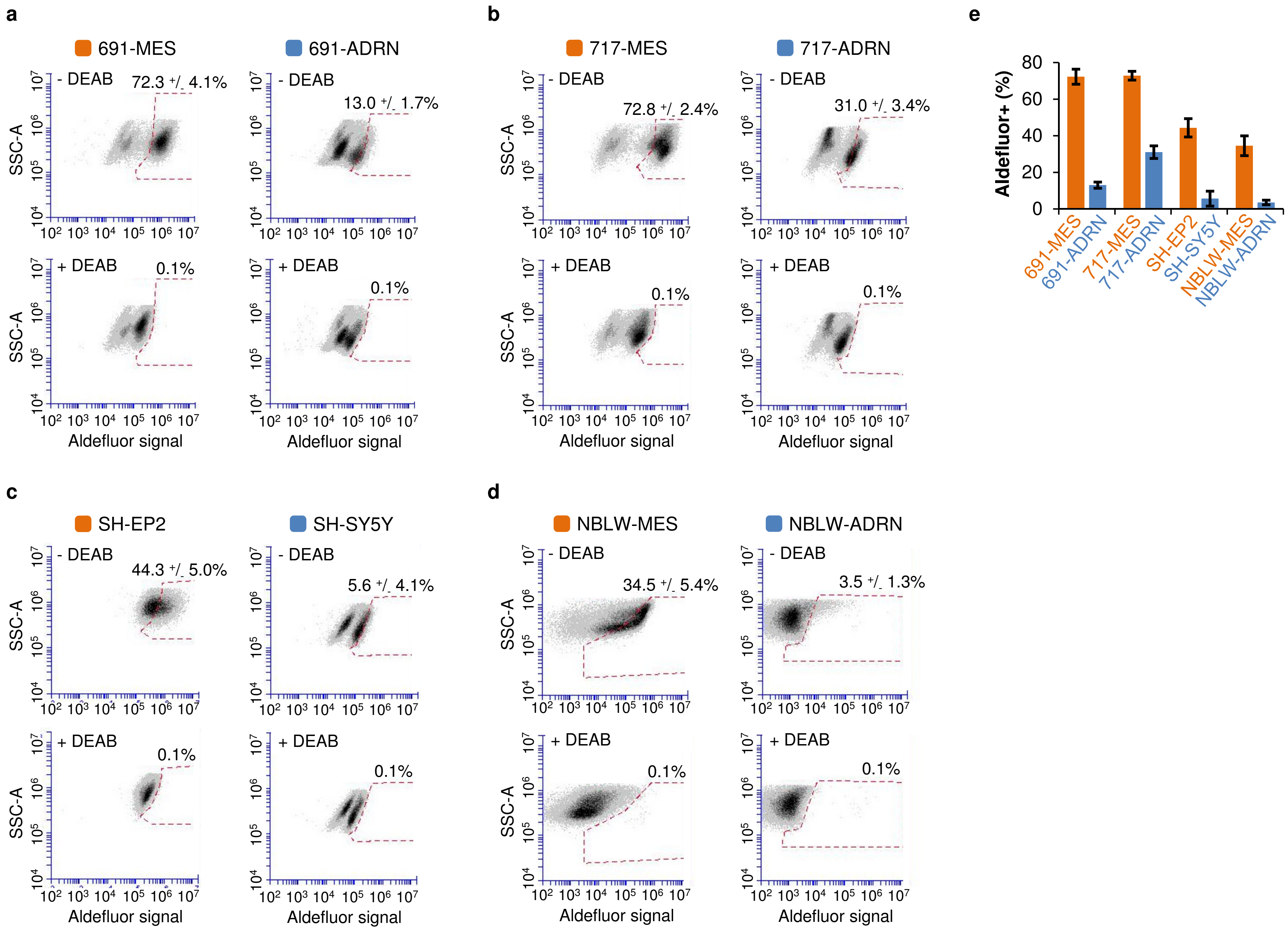

**Supplementary Figure 3 – related to Figure 1.**

**ALDH activity marks MES-type neuroblastoma cells.**

A-D. Flow cytometry analysis of ALDH activity measured by Aldefluor-assay in four isogenic MES and ADRN cell line pairs (A) 691-MES/691-ADRN, (B) 717-MES/717-ADRN, (C) SH-EP2/SH-SY5Y and (D) NBLW-MES/NBLW-ADRN in the absence (-, upper panel) or presence (+, lower panel) of the ALDH-inhibitor DEAB. The gate includes Aldefluor$^{positive}$ cells sensitive to DEAB-treatment. A representative FACS analysis from three independent Aldefluor experiments is shown. The percentages of Aldefluor$^{positive}$ cells are averages of three independent measurements +/- standard deviation. Aldelfuor signal is shown on the x-axis, side-scatter (SSC-A) is shown on the y-axis.

E. Summary of Aldefluor$^{positive}$ cells in the four isogenic MES and ADRN cell line pairs. Shown are the average measurements of three independent FACS experiments. In each experiment, 10,000-20,000 cells were analysed. The error bar indicates standard deviation. MES cell lines are shown in orange, ADRN cell lines are shown in blue. The percentage of Aldefluor$^{positive}$ cells is shown on the y-axis. Source data are provided as a Source Data File.

#### Supplementary Figure 4

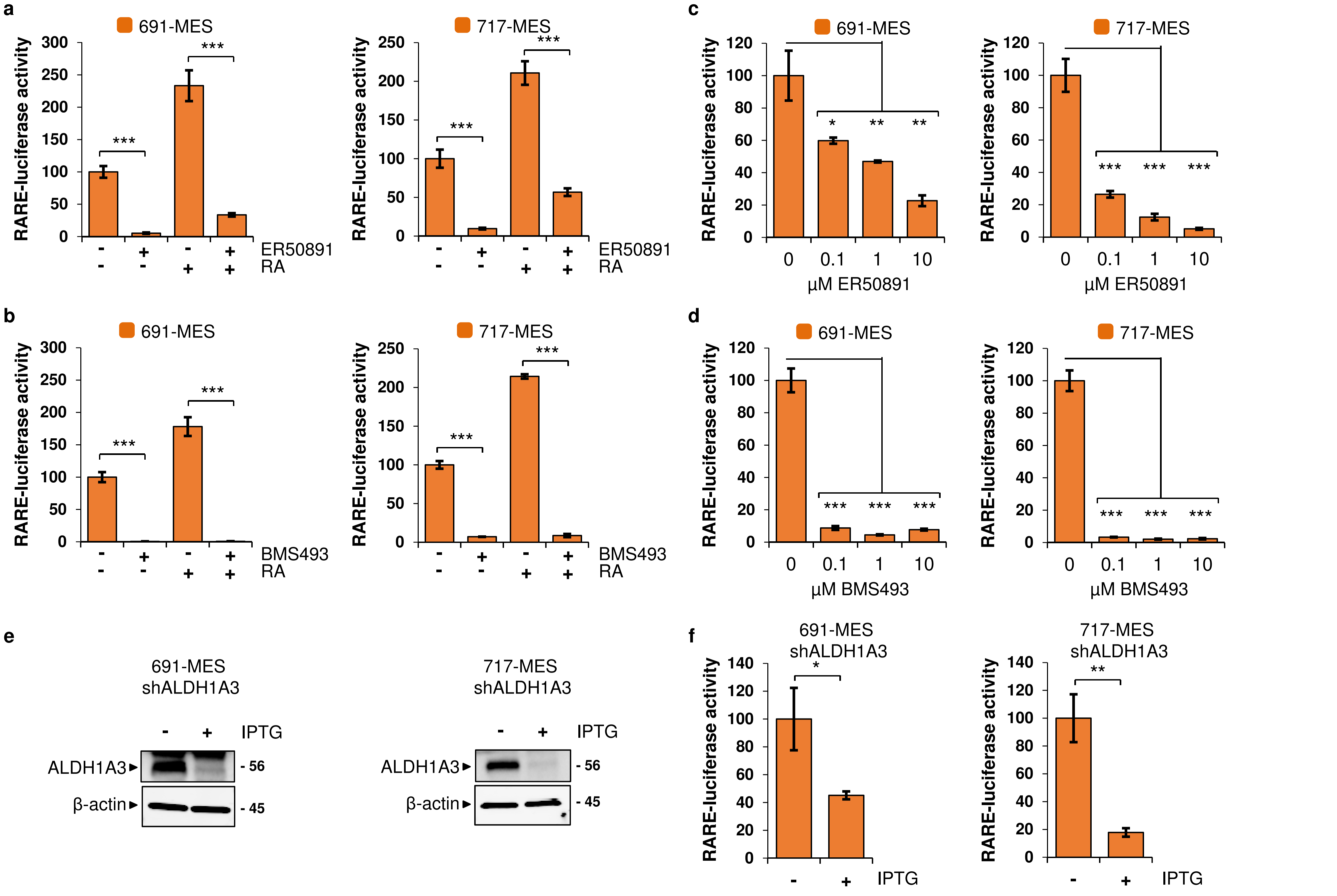

**Supplementary Figure 4 – related to Figure 1.**

**Inhibition of a RARE-reporter gene with endogenous activity in MES cells.**

A, B. RA reporter assay using a 3xRARE-luciferase construct in 691-MES and 717-MES cells. Cells were incubated for T=24 hours in the presence (+) or absence (-) of 100 nM RA in combination with 10 μM of the RARα inhibitor ER50891 (A) or 10 μM of the pan-RAR inhibitor BMS493 (B).

C, D. RA reporter assay (3xRARE-luciferase reporter gene) in 691-MES or 717-MES cells that were cultured in increasing concentrations of (C) the RARα inhibitor ER50891 or (D) the pan-RAR inhibitor BMS493.

E. Western blot analysis of ALDH1A3 in 691-MES (left) and in 717-MES (right) cells with IPTG-inducible shRNA targeting *ALDH1A3*. β-actin is shown as loading control. Molecular weight (in kDa) is indicated on the right.

F. RA reporter assay (3xRARE-luciferase reporter gene) in 691-MES and 717-MES cells with inducible shRNA targeting *ALDH1A3*. The normalized luciferase activities in A-D and G-H are ratios between firefly-luciferase values of the 3xRARE reporter and renilla-luciferase values of the transfection control. Error bars denote standard deviation. Two-sided Student's *t*-test assuming equal variance was used to calculate statistical significance, * $p < 0.05$, ** $p < 0.01$, *** $p < 0.001$. Source data for A-F are provided as a Source Data file.

#### Supplementary Figure 5

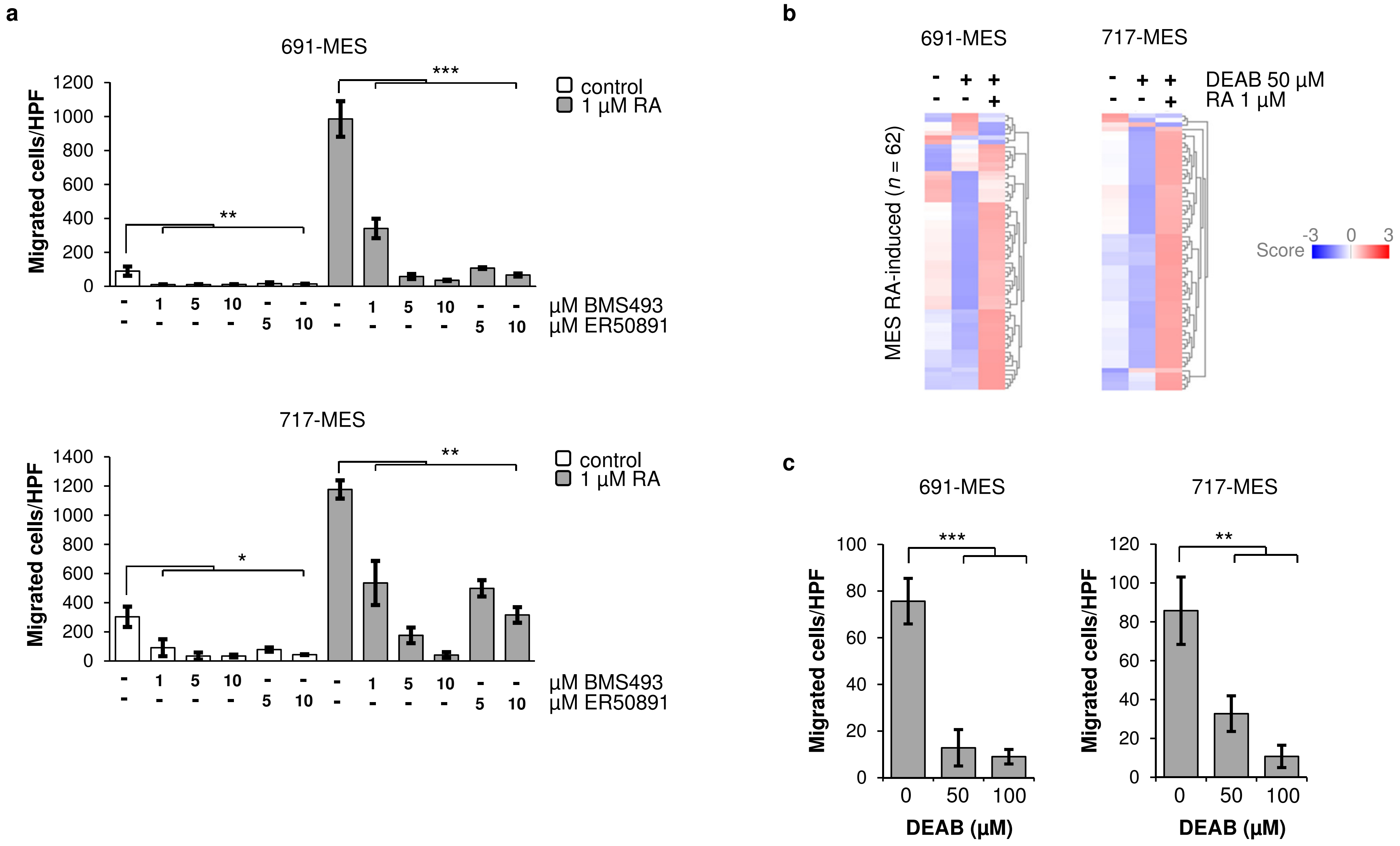

**Supplementary Figure 5 – related to Figure 2.**

**ALDH-activity and RA induce a motility programme in MES cells.**

A. Transwell migration assay of 691-MES and 717-MES cells in the absence (white) or presence (grey) of 1 μM RA in combination with increasing concentrations of the RARα inhibitor ER50891 or the pan-RAR inhibitor BMS493. Cells were allowed to migrate for 48 hours. Source data are provided as a Source Data File.

B. Heatmap visualization of the MES-RA$^{induced}$ gene signature in 691-MES (left) or 717-MES cells (right) treated for 72 hours with 50 μM DEAB or DMSO with or without 1 μM RA and analyzed by Affymetrix mRNA profiling. The list of RA-target genes in MES cells is available from Supplementary Table 2.

C. Cell migration assay of 691-MES and 717-MES cells treated with the ALDH-inhibitor DEAB. Cells were pre-incubated with DEAB for 4 days prior to seeding in a Boyden chamber to assess migration. Error bars in A and C denote standard deviation. Two-sided Student's *t*-test assuming equal variance was used to calculate statistical significance, * $p < 0.05$, ** $p < 0.01$, *** $p < 0.001$. Source data are provided as a Source Data File.

**Supplementary Figure 6**

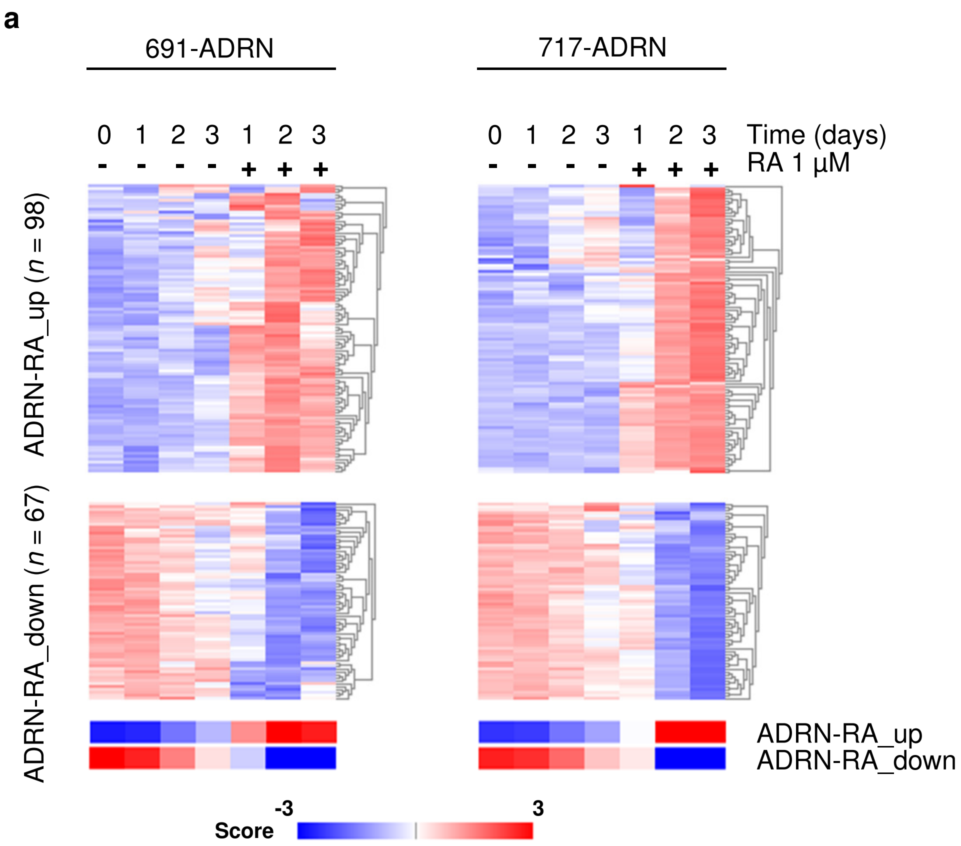


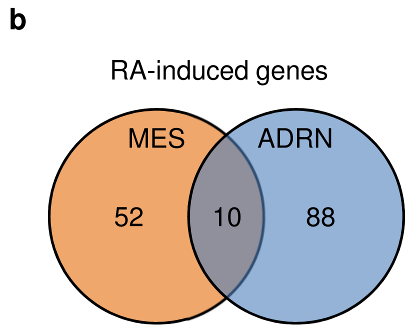

**Supplementary Figure 6 – related to Figure 2.**

**RA induces distinct gene-sets in MES and ADRN cells.**

A. Heatmap visualization of genes regulated by RA in ADRN cells. Expression is shown in 691-ADRN and 717-ADRN cells cultured for indicated time points in the presence (+) or absence (-) of 1 μM RA. Cumulative signature scores for both up- and down-regulated RA responsive gene-sets are indicated at the bottom for each time point. Time is indicated in days. The list of RA-target genes in ADRN cells is available from Supplementary Table 2.

B. Venn diagram depicting the overlap of MES-specific RA-induced genes (ROL-RA responsive in MES cell lines 691-MES and 717-MES) and ADRN-specific RA-induced genes (RA responsive in ADRN cell lines 691-ADRN and 717-ADRN).

### Supplementary Figure 7

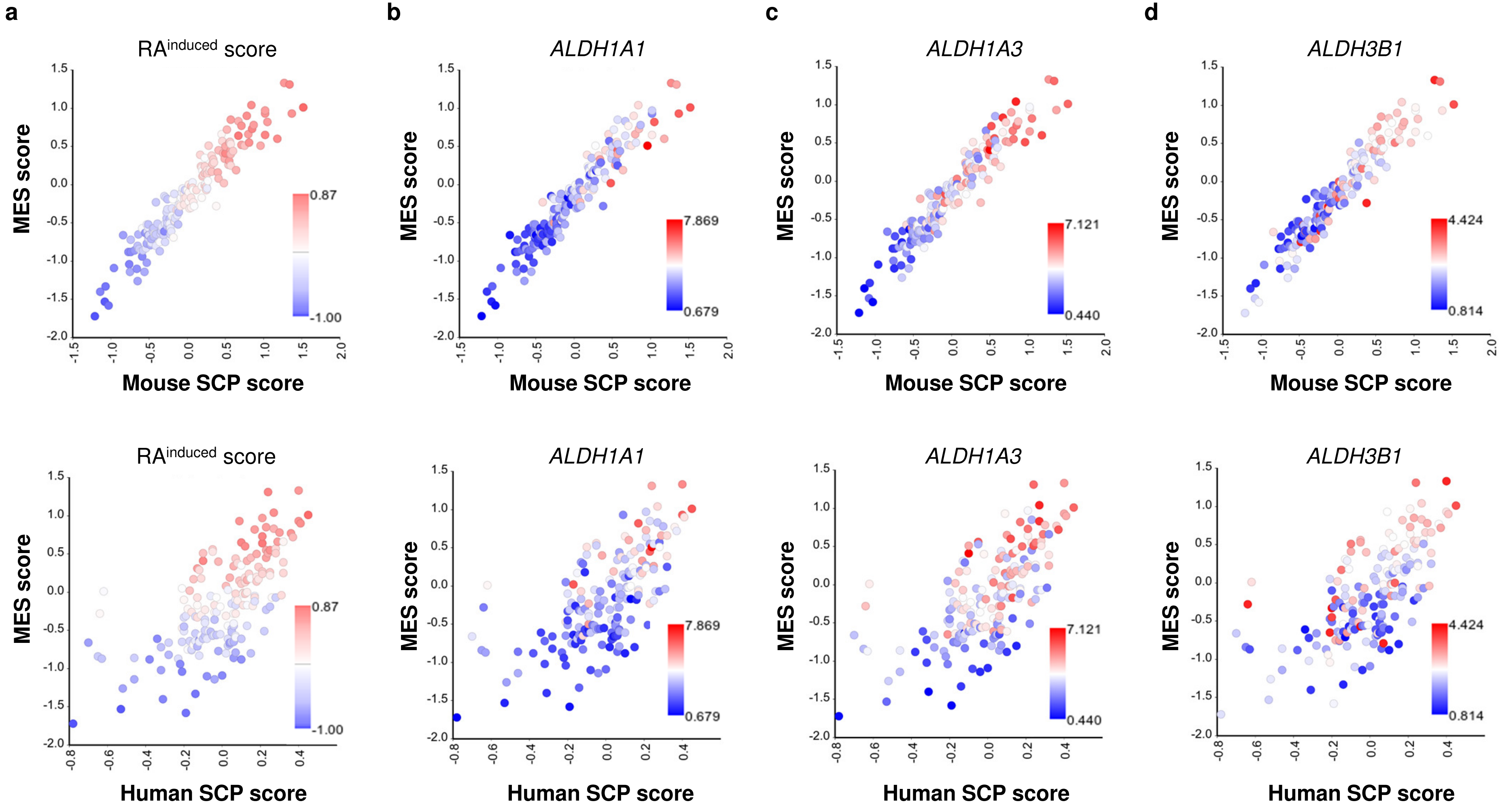

**Supplementary Figure 7 – related to Figure 4.**

**Analysis of RA gene signature scores in human neuroblastoma.**

A. Analysis of mRNA signature scores for mouse SCP cells (x-axis) and human neuroblastoma MES cells (y-axis) in a panel of INSS stage 4 human neuroblastoma tumors (n=183). Each tumor is represented by a dot. A signature score of the $\text{RA}^{\text{induced}}$ target genes in MES cells is visualized for each tumor. The signature scores represent the summed z-scores of genes in the signature.

B-D. mRNA expression ($^{2}$log-transformed) of (B) *ALDH1A1*, (C) *ALDH1A3*, (D) *ALDH3B1* on the INSS stage 4 neuroblastoma shown in panel A.

E. Analysis of gene expression signature scores for human SCP cells (x-axis) and for human neuroblastoma MES cells (y-axis) in a series of 183 INSS stage 4 neuroblastoma, similar to the analysis in A.

F-H. mRNA expression ($^{2}$log-transformed) of (F) *ALDH1A1*, (G) *ALDH1A3*, (H) *ALDH3B1* on the INSS stage 4 neuroblastoma shown in panel E.

**Supplementary Figure 8**

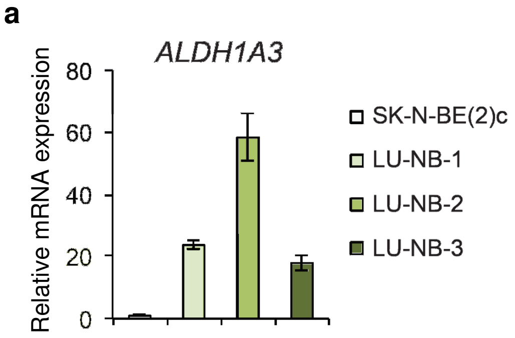


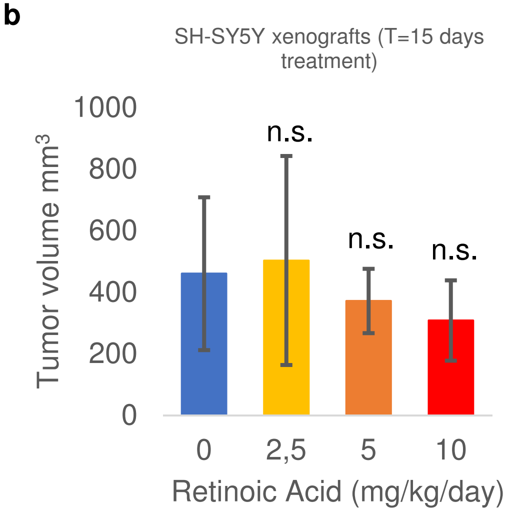


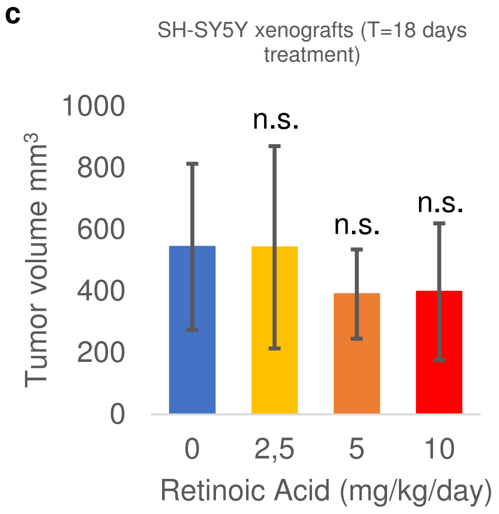


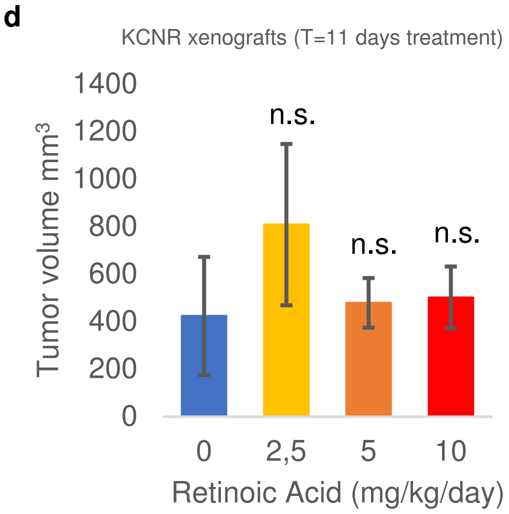


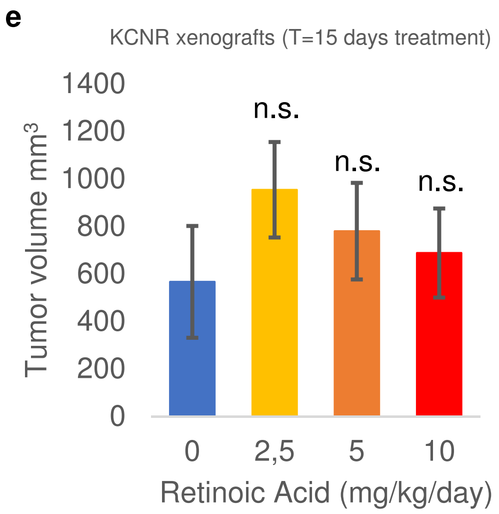

**Supplementary Figure 8 – related to Figure 4.**

**Analysis of *ALDH1A3* expression and RA-treatment *in vivo*.**

A. mRNA expression of *ALDH1A3* in cells from three neuroblastoma PDX models (LU-NB-1, LU-NB-2, and LU-NB-3) and in the neuroblastoma cell line SK-N-BE(2)c.

B-D. Retinoic acid treatment of neuroblastoma xenografts. SH-SY5Y (B, C) and KCNR (D, E) xenografts were treated with 0, 2.5, 5 or 10 mg/kg body weight for 5 consecutive days, followed by 2 days off treatment for each week during the experiment. Average tumor volume (n=3 mice per treatment group) is plotted for SH-SY5Y at T=15 and T=18 (B, C) as well as for KCNR at T=11 and T=15 (D, E) days after start of treatment. Source data of B-D are provided as a Source Data File.
